# Quantitative MRI provides markers of intra-, inter-regional, and age-related differences in young adult cortical microstructure

**DOI:** 10.1101/139568

**Authors:** Daniel Carey, Francesco Caprini, Micah Allen, Antoine Lutti, Nikolaus Weiskopf, Geraint Rees, Martina F. Callaghan, Frederic Dick

## Abstract

Measuring the structural composition of the cortex is critical to understanding typical development, yet few investigations in humans have charted markers in vivo that are sensitive to tissue microstructural attributes. Here, we used a well-validated quantitative MR protocol to measure four parameters (R_1_, MT, R_2_*, PD*) that differ in their sensitivity to facets of the tissue microstructural environment (R_1_, MT: myelin, macromolecular content; R2*: paramagnetic ions, i.e., iron; PD*: free water content). Mapping these parameters across cortical regions in a young adult cohort (18-30 years, N=93) revealed expected patterns of increased macromolecular content as well as reduced tissue water content in primary and primary adjacent cortical regions. Mapping across cortical depth within regions showed decreased expression of myelin and related processes – but increased tissue water content – when progressing from the grey/white to the grey/pial boundary, in all regions. Charting developmental change in cortical microstructure, we found that parameters with the greatest sensitivity to tissue myelin (R_1_ & MT) showed linear increases with age across frontal and parietal cortex (change 0.5-1.0% per year). Overlap of robust age effects for both parameters emerged in left inferior frontal, right parietal and bilateral pre-central regions. Our findings afford an improved understanding of ontogeny in early adulthood and offer normative quantitative MR data for inter- and intra-cortical composition, which may be used as benchmarks in further studies.

**Highlights:**

1. We mapped multi-parameter maps (MPMs) across and within cortical regions
2. We charted age effects on myelin and related processes at mid-cortical depth
3. Inter- and intra-regional differences in MPMs emerged at primary and association cortex
4. Iron-sensitive R_2_* map foci tended to overlap MPMs sensitive to myelin (R_1_, MT)
5. R_1_ and MT increased with age (0.5-1.0% per year) in frontal and parietal cortex

---

A core challenge for human neuroscience is the design of robust anatomical imaging methods that are sensitive to inter-regional differences in tissue properties, and to profiles of intra-cortical tissue change from the grey-white border to the pial surface in any one region. The parcellation of human cortex based on cyto- and myeloarchitectonic boundaries has been a major pursuit since the work of Brodmann and Flechsig in the early 20th century (Sereno et al., 2013; Glasser et al., 2016; Turner, 2015; Nieuwenhuys, 2013; Nieuwenhuys et al., 2014; Zilles et al., 2015). However, it is only recently that such questions have been addressed *in-vivo* in humans. This is made possible by the use of magnetic resonance imaging (MRI), which provides data for morphometry (Ashburner & Friston 2000; Dale et al., 1999; Fischl et al., 1999a, 1999b) or microstructure (Weiskopf et al., 2015).

The MR signal is sensitive to many important tissue properties, such as iron content, myelin, cell density and water content; however, the contrast-weighted images (T_1w_, T_2w_) typically used in MRI reflect a complex mix of these properties that can vary non-linearly across the imaged volume. By comparison, *Quantitative MRI* (Bock et al., 2013; Barazany & Assaf, 2012; Dinse et al., 2013, 2015; Stüber et al., 2014; Marques et al., 2010; for review, see Turner, 2015, 2016; Bazin et al., 2014; Cohen-Adad, 2014; Sereno et al., 2013; Dick et al., 2012) can be used to map specific MRI properties of tissue in order to provide indices of microstructure, myelination and related cellular processes (Helms et al., 2008a, 2009; Weiskopf et al., 2013; Lutti et al., 2014) in a time-efficient manner with high spatial specificity. It thus provides the opportunity to acquire a multi-modal, whole-brain view of developmental changes in underlying tissue properties.

In the multi-parameter mapping (MPM) quantitative imaging protocol (Weiskopf et al., 2013; Callaghan et al., 2014b; Helms et al., 2008a, 2008b; Lutti et al., 2014), multiple maps are constructed to probe different tissue attributes. These are 1) the longitudinal relaxation rate, R_1_ = 1/T_1_ (sensitive to myelin, macromolecular content and water); 2) the effective transverse relaxation rate, R_2_* = 1/T_2_* (sensitive to susceptibility effects due to paramagnetic ions, most notably iron); 3) Magnetisation Transfer (MT; sensitive to myelin, macromolecular content and bound water fraction); and 4) effective Proton Density (PD*; sensitive to free water content and residual R_2_* related effects) (Weiskopf et al., 2011; Callaghan et al., 2014a, 2014b; Lutti et al., 2014; Stüber et al., 2014; Fukunaga et al., 2010). These methods allow quantitative measurement of inter- and intra-regional differences in tissue properties (e.g., Cohen-Adad et al., 2012; 2014; Govindarajan et al., 2015; Dinse et al., 2015) including age-related changes in subcortical fibre tract myelination (Yeatman et al., 2014), pathological changes in neurotrauma (Freund et al., 2013), maturation effects (Whitaker et al., 2016), and age-related tissue de-myelination (Callaghan et al., 2014a), whilst affording the means to do so in relation to functional ability (e.g., Gomez et al., 2017). Such myelin mapping methods have also been used to identify the heavily-myelinated boundaries of visual (Sereno et al., 2013; but see Abdollahi et al., 2014), primary auditory (Dick et al., 2012; de Martino et al., 2014; Sigalovsky et al., 2006), and somatomotor areas (Carey et al., 2017), when relating these regions to function.

Charting the normal development and aging of human cortical tissue is a fundamental goal of neurobiology, and is also critical for accurately characterizing atypical development, individual differences, and short- and long-term plasticity. Development is reported to follow a posterior-to-anterior gradient with primary areas maturing earliest in life and association areas, which mediate higher-order functions, developing later (Gogtay et al., 2004; for review of processes, see Marsh et al., 2008). At earlier points in development through adolescence, there is evidence to suggest that deviation from typical trajectories may increase vulnerability to psychiatric disorders (Thompson et al., 2001; Greenstein et al., 2006; Sowell et al., 2003; Shaw et al., 2007) whereas in later life, such deviation may be indicative of neurodegenerative decline, for which age is often the greatest predictor (Barkhof et al., 2009; Bartzokis, 2004; Bartzokis, 2011; Frisoni et al., 2010). A cornerstone in the development of mature cortex is the emergence of myelinated fibres within the cortical sheet (Flechsig, 1920; Yakovlev & Lecours, 1967; Deoni et al., 2015). Though the exact trajectories are unclear, the rate at which change occurs – and the age at which development stabilizes – are thought to be region-specific (e.g. Yeatman et al., 2014; Whitaker et al., 2016) and to interact with functional organization (e.g. Yeatman et al., 2012; Gomez et al., 2017).

To date, few quantitative imaging studies have explored developmental changes in tissue composition across cortex from late adolescence to the mid thirties. This is a crucial age range to characterize, not least because it is the 'sample of choice' for the vast majority of structural and functional MRI studies. Here, we used the MPM protocol (Weiskopf et al., 2013; Callaghan et al., 2014a, 2014b; Helms et al., 2008a, 2008b) to explore potential parameter-specific (R_1_, MT, PD*, R_2_*) variation in tissue over the depth of the cortical sheet, and across a range of cortical regions. Further, we charted age-related differences in cortical microarchitecture across early adulthood. We mapped a set of normative, cortical-depth-specific regional MPM values for young adults that can be used as reference values for future studies. Moreover, we found considerable, region-specific age-related changes in parameters related to the degree of tissue myelin content and myelin-related processes.

## Materials and Methods

### 2.1 Participants

Participants were 93 right-handed healthy adults (mean age ± SD: 23.6 ± 4.3; range: 18-39; 57 female, 36 male). The study received approval from the local ethics committee. All scanning took place at the Wellcome Trust Centre for Neuroimaging (WTCN), London.

Participants were sampled over approximately 24 months. Thirty-four participants were recruited as part of a study of musicianship and consisted of expert violinists (*n* = 18; mean age ± SD: 22.8 ± 2.8; 13 female, 5 male) and closely matched non-musicians (*n* = 16; mean age ± SD: 23.3 ± 3.1; 12 female, 4 male). All had completed or were enrolled in a university degree, and were recruited from the University of London, music conservatories in London, and local participant pools. We analyzed data for effects of violin expertise and will report these findings in a subsequent report. In brief, effects of violin expertise in cortex were modest and emerged only in ROI analyses of primary auditory cortex, where we found limited evidence of significant age-related effects in the present study.

The remaining participants (*n* = 59; mean age ± SD: 23.9 ± 4.9; 32 female, 27 male) were sampled from the general population through local participant pools. These subjects took part in three experiments: one exploring the potential association between auditory perceptual abilities, musicianship, tonotopic organization and structural properties of the auditory cortex (data not reported here), one investigating the relationship of trait empathy and brain microstructure (Allen, Frank, et al., 2017), and a third investigating metacognition and MPM assays (Allen et al., 2017).

### 2.2 Data Acquisition

The multi-parameter mapping protocol data (Weiskopf et al., 2013; Lutti et al., 2010, 2012) were acquired at the WTCN using a 3T whole-body Tim Trio system (Siemens Healthcare) with radiofrequency body coil for transmission and a 32- channel head coil for signal reception. The MPM protocol consisted of three differently weighted 3D multi-echo FLASH acquisitions acquired with 800 micron isotropic resolution. Volumes were acquired with magnetization transfer (MT_w_), T_1_-(T_1w_), and proton density (PD_w_) weighting. The MT weighting was achieved through application of a Gaussian RF pulse (4ms duration, 220° nominal flip angle) applied 2kHz off-resonance prior to non-selective excitation.

Two further scans were collected to estimate participant-specific inhomogeneities in the RF transmit field (B_1_^+^) using a 3D EPI acquisition of spin-echo (SE) and stimulated echo (STE) images as described in Lutti et al. (2010) (slice thickness: 4 mm; matrix size: 64 × 48 × 48; field-of-view: 256 × 192 × 192 mm^3^; bandwidth: 2298 Hz/pixel; SE/STE acquisition time post-excitation: 39.38 ms/72.62 ms; TR: 500 ms). In addition, a map of the B_0_ field was acquired and used to correct the B_1_+ map for off-resonance effects (Lutti et al., 2010; see also Weiskopf et al., 2006) (voxel size: 3 × 3 × 2 mm^3^; slice thickness: 4mm; field-of-view: 192 × 192 mm^2^; 64 slices, 1mm gap; bandwidth: 260 Hz/pixel; TE1 10 ms, TE2 12.46 ms; TR: 1020 ms; flip angle: 90°).

The sequence settings of the MPM protocol were modified following collection of data for the musicianship sample (cohort 1, *n* = 34), reflecting the on-going development of the MPM sequences at the WTCN. Cohort-specific details follow.

#### 2.2.1 Cohort 1

A field of view of 256 × 224 × 166 mm^3^ was used with a matrix size of 320 × 280 × 224. The PD_w_ and T_1w_ volumes were acquired with a TR of 25.25 ms and a flip angle of 5° and 29° respectively. The MT_w_ volume was acquired with a TR of 29.25 ms and excitation flip angle of 9°. The excitation employed a hard pulse; RF spoiling was used with an increment of 50 degrees, and gradient spoiling producing 6*pi dephasing across a voxel. The acquisition was accelerated by using GRAPPA (with a parallel imaging factor of 2 with 18 integrated reference lines) in the phase-encoded direction (AP) and by a partial Fourier acquisition in the partition direction (RL, with factor 6/8). To improve image quality (maximize SNR and minimize geometric distortion at the same time), eight gradient echoes with alternating readout polarity were acquired with high readout bandwidth (460 Hz/pixel) with echo times ranging from 2.39 ms to 18.91 ms in steps of 2.36 ms.

A fixed modification of MPM image slab orientations (30 degrees) was applied for some subjects to counter image artifact due to eye movement and blinking; this change in the acquisition did not yield any significant differences (at *p* < 0.01, whole-brain uncorrected) for any map (R_1_, PD, MT, R_2_*) between subjects with and without slab rotation (see footnote 1). Image slabs for field maps were all non-rotated along the axial orientation.

The B_1_ mapping acquisition consisted of 15 measurements with nominal flip angle ranging from 135° to 65° in 5° decrements. The total scanning time of the MPM protocol was approximately 37 minutes.

#### 2.2.2 Cohort 2

For the second cohort, the MPM protocol was modified to improve accuracy by accounting for non-linearities in the transmit chain (Lutti and Weiskopf, 2013). To achieve this, different flip angles for the PD-, MT- (both 6°) and the T_1_- (21°) weighted acquisitions were achieved by scaling the duration of the pulse while maintaining a constant B_1+_ amplitude (i.e. a consistent operating point for the RF amplifier) that additionally matched that used for the B_1_+ mapping sequence. Gradient echoes were again acquired with alternating readout gradient polarity using a readout bandwidth of 488Hz/pixel. Eight equidistant echo times ranging from 2.34 to 18.44 ms in steps of 2.3 ms were acquired for the PD_w_ and T_1w_ acquisitions. Only the first six echoes were acquired for the MT-weighted acquisition in order to maintain a 25 ms TR for all of the FLASH volumes. To further accelerate the data acquisition, the partial Fourier acquisition scheme in the partition direction was replaced by parallel imaging with an acceleration factor of 2 again using the GRAPPA algorithm, now with 40 integrated reference lines in each phase-encoded direction. A 30 degree slab rotation was used for all acquisitions in this cohort.

The B_1_ mapping acquisition consisted of 11 measurements with nominal flip angle ranging from 115° to 65° in 5° decrements. The total scanning time of the MPM protocol was approximately 26 minutes.

### 2.3 Procedure

Participants provided written informed consent and were screened for contraindications for MRI. B_1_+ and B_0_ field maps were collected at the beginning of each session, followed by the MT, PD_w_, and T_1w_ scans. Participants’ eye and head movements were monitored using an eye tracker (Eyelink 1000 Core System) during scanning runs. Rest breaks of several minutes were provided between scans as required.

#### 2.3.1 Data Pre-Processing

Images were pre-processed using the Voxel Based Quantification (VBQ) toolbox in SPM 8. In brief, regression of the log signal from the echoes of all weighted volumes were used to calculate a map of R_2_* using the ordinary least squares ESTATICS approach (Weiskopf et al., 2014). The set of echoes for each acquired weighting were then averaged to increase the signal-to-noise ratio (Helms and Dechent, 2009). This was done using only the first six echoes for Cohort 2. The 3 resulting volumes were used to calculate MT, R_1_, and PD* maps as described in Helms et al. (2008a, 2008b) and Weiskopf et al. (2013). Quantitative R_1_ values at each voxel were estimated based on the rational approximation of the Ernst equation described by Helms et al. (2008a). To maximize the accuracy of the R_1_ map, these maps were corrected for transmit field inhomogeneities by constructing a map from the calibration data according to the procedure detailed in Lutti et al. (2012). The R_1_ maps were also corrected for imperfect spoiling characteristics using the approach described by Preibisch and Deichmann (2009). The MT map was constructed using the procedure described in Helms et al. (2008b). This is a semi-quantitative metric depicting the percentage loss of magnetization resulting from the MT pre-pulse used and differs from the commonly used MT ratio (percentage reduction in steady state signal) by explicitly accounting for spatially varying T_1_ relaxation times and flip angles (Weiskopf et al., 2013). Finally, PD* maps were estimated from the signal amplitude maps by adjusting for receive sensitivity differences using a post-processing method similar to UNICORT (Weiskopf et al., 2011). To make the PD*maps comparable across participants, they were scaled to ensure that the mean white matter PD* for each subject agreed with the published level of 69% (Tofts, 2003). This quantity is referred to as effective PD (PD*) because it was calculated based on the average FLASH volumes and there was no correction for R_2_* signal decay.

Following reconstruction of multi-parameter images, all images were manually inspected for any evidence of alignment difficulties, head movement or other image artifacts (e.g., aliasing) by a rater who was blind to subject identity.

#### 2.3.2 Cortical Surface Reconstruction

Participants’ cortical surfaces were reconstructed using FreeSurfer (v. 5.3; Dale et al., 1999). Use of multi-parameter maps as input to FreeSurfer can lead to localized tissue segmentation failures due to boundaries between the pial surface, dura matter and CSF showing different contrast compared to that assumed within FreeSurfer algorithms (Lutti et al., 2014). Therefore, an in-house FreeSurfer surface reconstruction procedure was developed to overcome these issues.

Full details of the processing pipeline are provided in supplemental methods."

### 2.4 Data analyses

Following cortical surface reconstruction, R_1_, MT, R_2_* and PD* data were mapped onto participants’ cortical surfaces in FreeSurfer. Whole-brain vertex-wise analyses were subsequently performed. Description follows below (2.4.1 & 2.4.2).

#### 2.4.1 MPM data extraction

First, all input volumes were re-sampled to a finer image resolution (0.6 mm^3^ isotropic) and all subjects were rotated to the same (canonical) orientation, using the AFNI 3dwarp routine (-deoblique flag). MPM data were then mapped onto each subject’s surface (using the FreeSurfer mri_vol2surf routine). For each reconstructed hemisphere, quantitative data were sampled along the normal to each surface vertex, for cortical depth fractions from 0.1 (i.e., above white matter surface boundary) to 0.9 (i.e., beneath pial surface boundary) in increments of 0.1 (see Dick et al., 2012).

#### 2.4.2 Analyses

For each relaxation parameter (R_1_, MT, R_2_*, PD*), we first created cross-subject hemisphere-wise average maps for each cortical depth sampling fraction (0.1-0.9) using cortical-surface-based methods with curvature-based alignment (Fischl et al., 1999; Hagler & Sereno, 2006; Dick et al., 2012; Sereno et al., 2013). In each of a series of cortical regions-of-interest, we extracted hemisphere-wise mean estimates of each relaxation parameter across depth fractions. Cortical regions were defined from a standard FreeSurfer atlas (aparc.2009), sampled onto each subject’s cortical surface during reconstruction (regions were: superior pre-central sulcus; subcentral gyrus/sulcus; inferior pre-central sulcus; inferior frontal sulcus; middle frontal sulcus; superior parietal gyrus; angular gyrus; Heschl’s gyrus; inferior occipital gyrus/sulcus; superior temporal gyrus/planum temporale; probabilistic area MT; probabilistic V1; posterior collateral sulcus; superior occipital gyrus; parieto-occipital sulcus; subparietal sulcus; middle cingulate gyrus/sulcus). ROI mean estimates for each relaxation parameter were averaged across hemispheres at each cortical depth fraction sampled (0.1-0.9).

For age-based analyses, we used participants’ age in whole years as a continuous linear regressor at each vertex per hemisphere, with initial analyses carried out in Qdec (FreeSurfer v. 5.3). For age analyses, each subject’s data were smoothed with a surface (2D) kernel of 10mm FWHM. Unless otherwise specified, age analyses are reported sampling at 0.5 cortical depth, that being the most representative of mid-cortical depth profiles (Dick et al., 2012; Sereno et al., 2013; Lutti et al., 2014; cf. Waehnert et al., 2014).

Previous studies have found that R_1_ values are associated with variation in the local curvature and thickness of the cortex; thus, R_1_ measurements tend to be increased in thicker, more highly convex regions (e.g., gyral crowns) (Sereno et al., 2013; Dick et al., 2012; Waehnert et al., 2014). In addition, changes in the MPM acquisition protocol between the cohorts we scanned here (see 2.2.1 & 2.2.2) were associated with differences in MPM map values. The second cohort, scanned with the protocol that better addressed non-linearities in the transmit chain, showed consistently greater (up to ∼15%) R_1_ values with a gently spatially varying pattern that could be reproduced by comparing results in a single individual subject scanned across both protocols. Therefore, to control for these effects, we regressed out local curvature, cortical thickness and MPM cohort (see also Grydeland et al., 2013); we then performed Pearson correlations at each vertex between age and MPM values that were residualized by curvature, thickness and MPM-cohort. In general, curvature-, thickness-, and cohort-residualized age regressions were similar to age regressions using raw parameter values.

To produce unbiased estimates of age effects on each parameter we used a ‘leave-one-out’ jackknife procedure. Here, we first performed vertex-wise age-MPM Pearson correlations for the full cohort, and then repeated the procedure iteratively omitting one subject in each instance. Pearson *r*-values for the full cohort and each leave-one-out partial estimate were Fisher z-transformed, and a mean of partial estimates calculated. Jackknife estimates at each vertex were calculated as: (N)(T) - (N-1)(Tm), where N = 93; T = full cohort z-transformed *r*-value; Tm = mean of partial estimates (z-transformed before averaging). Finally, vertex-wise Jackknife estimates were re-transformed to *r*-values and corresponding *p* values were calculated. The jackknifing procedure was performed in regression models with age as a vertex-wise predictor of raw MPM values, and also in regression models with age as a vertex-wise predictor of cohort-, thickness- and curvature- residualized maps.

Jackknifed statistical maps were thresholded using peak-level False-Discovery Rate (FDR) correction (Benjamini & Hochberg, 1995); FDR-corrected *q* < 0.05, per hemisphere. For illustrative purposes, we also identified regions where significant jackknifed effects of age overlapped for both the R_1_ and MT multi-parameter maps. Note that these maps show greatest sensitivity to cortical myelin, and thus were of central interest here (see Lutti et al., 2014; Sereno et al., 2013). Per hemisphere, we determined the vertices that survived FDR-correction (*q* < 0.05) for jackknifed age analyses of the cohort-, thickness- and curvature-residualized MPMs (i.e., R_1_ & MT). Using Matlab, we then created a binary mask per hemisphere corresponding with vertices where the jackknifed model results for both MPMs showed FDR-significant effects of age. Clusters of vertices reflecting the overlap of the age effects for the two MPMs (R_1_ & MT) were extracted and defined as ROIs on a standard cortical surface; ROIs were sampled onto each subject’s cortical surface. Across each of these ROIs, we plotted the linear relationship between age and subject-wise ROI means for R_1_ and MT.

## Results

Here, we explored intra- and inter-regional differences in quantitative markers of tissue microstructure, using a multi-parameter mapping protocol that affords quantitative MR proxies for myelin-related tissue processes. Further, we used a cross-sectional design to explore the effects of age across early adulthood on cortical myelination. To develop a clearer understanding of inter- and intra-regional myelin-related tissue properties, we produced average maps for each of the MPMs across the cortical sheet, sampling halfway through cortex; moreover, we charted differences in MPM parameters over the depth of the cortical sheet, within and between a range of cortical ROIs.

### 3.1 Group average MPM results

Average MPMs (Fig. 1) revealed the expected inter-regional differences in cortical myelin and myelin-related processes, in line with previous literature. Parameters that show greatest sensitivity to myelin and related processes (R_1_, MT, and R_2_*), had highest values within motor and sensory regions, including somatomotor, auditory, and visual cortex. Further to previous studies (Glasser & Van Essen, 2011; Glasser et al., 2016; Sereno et al., 2013; Waehnert et al., 2016; Bock et al., 2013), R_1_, MT and R_2_* revealed strips of dense myelination over pre-central and post-central regions, reflecting areas 4 and 3b/1, respectively, with an intervening lower-myelin septum (likely area 3a; Geyer, 2013; Dinse et al., 2015; Stüber et al., 2014; see also Glasser & Van Essen, 2011). R_1_, MT and R_2_* also revealed the heavily myelinated auditory core at the most medial aspect of Heschl’s gyrus (Dick et al., 2012; Sigalovsky et al., 2006; de Martino et al., 2015); planum temporale and parts of lateral superior temporal gyrus (STG) additionally showed elevated R_1_ and MT values (see Glasser & Van Essen, 2011; Glasser et al., 2016; Sigalovsky et al., 2006). In visual areas, R_1_ and R_2_* exposed the heavily myelinated V1 extending across the calcarine sulcus (Sereno et al., 2013; Fracasso et al., 2016; see also Cohen-Adad et al., 2012); however, parameter MT showed high values that were restricted to gyral banks flanking the calcarine sulcus (Fig. 1, parameter MT medial surface panels). A possible source of this difference is the local contribution of myelin differences to macromolecular effects at gyri, versus the more anatomically diffuse effects of iron associated with oligodendrocyte cell bodies – contributing in part to the R_1_ signal, via R_2_* – found across sulci (e.g., Stüber et al., 2014; see 3.3, and discussion, 4.1). Alternatively, local patterns of cortical folding and curvature may have influenced the detection of macromolecular content by MT in highly concave cortical regions; cortical thickness in sulcal depths (and indeed much of V1) is roughly equal to the voxel dimensions, and therefore the contribution of the thin myelinated layers to the overall contrast will be attenuated.

**Figure 1.**
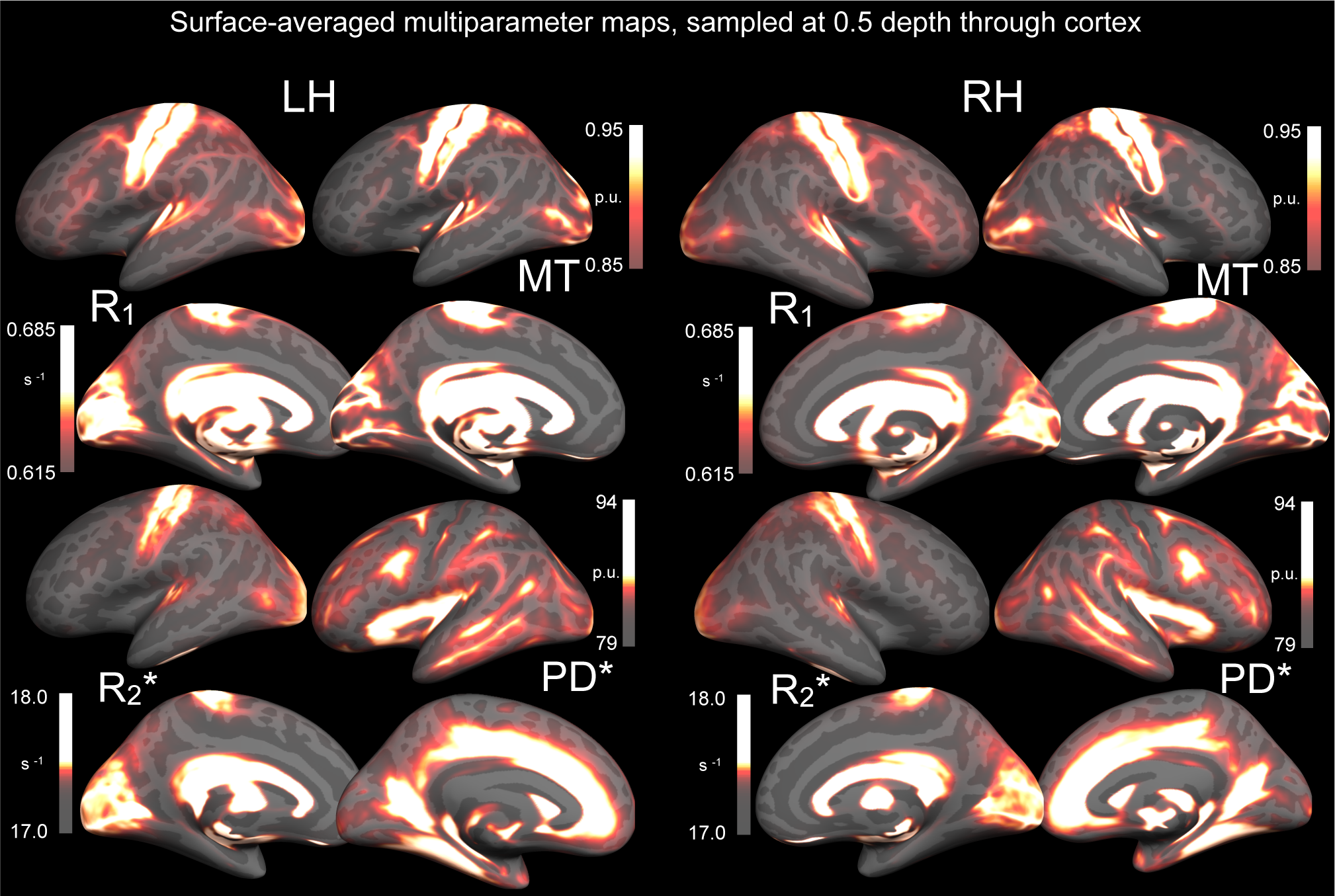
Surface-averaged multi-parameter maps for R_1_, MT, R_2_* and PD*. Maps reflect full cohort means, presented with all values sampled at 0.5 cortical depth. Parameter maps are shown for left and right hemisphere lateral and medial surfaces. Per hemisphere, leftmost column presents R_1_ (top) and R_2_* (bottom); rightmost column presents MT (top) and PD* (bottom). Heat scales present scale value range for surface overlay, with scale midpoints at centre. Note differing measurement units (R_1_ & R_2_*: s-1; MT & PD&: percentage units [PU]).

R_1_, MT and R_2_* additionally revealed higher visual areas including V6 (medial surface, dorsal to V1), V3/V3a (lateral surface, dorsal to V2), and area MT (proximal to postero-lateral bounds of inferior temporal sulcus). R_1_ and MT also revealed several heavily myelinated cortical regions posterior to post-central gyrus, likely including multi- modal VIP and LIP (Sereno et al., 2013; Glasser & Van Essen, 2011; Waehnert et al., 2016; Huang et al., 2012). Foci of high R_1_ and MT reflecting the frontal eye fields also emerged, lying proximal to the dorsal-most aspect of the middle frontal gyrus and the boundary with pre-central gyrus (see Glasser et al., 2016; Glasser & Van Essen, 2011). PD* revealed a more distinct patterning of regions than the other MPMs, reflecting its high affinity for unbound protons (i.e., tissue water; Baudrexel et al., 2016). Regions typically low in myelination (see Nieuwenhuys, 2013; Nieuwenhuys et al., 2014; Geyer, 2013; Zilles et al., 2015) tended to show highest PD* values, including: the circular sulcus and adjoining insular cortex; cingulate gyrus and sulcus; medial prefrontal cortex; collateral sulcus; parieto-occipital sulcus and regions anterior to V1; superior temporal sulcus; middle temporal gyrus; inferior frontal sulcus; and presumptive Area 3a.

### 3.2 Intra-regional MPM results

Across depth fractions, we observed the expected pattern of decrease in R_1_ and MT values over all cortical ROIs (Fig. 2a & 2b). Reduction in R_1_ and MT values broadly followed a decaying pattern, with sharpest decreases at depth fractions close to the white matter and pial surfaces (0.1 and 0.9, respectively; see Fig. 2a & 2b). In line with previous work (Annese et al., 2004; Dinse et al., 2015), this likely reflects the reduction in myelinated fibre density with progression away from the white matter surface toward the pial surface. In modeling the reduction in R_1_ and MT values over the cortical sheet in each ROI, we found that cubic trends provided the best fit in every region, versus linear and quadratic trends (fits achieved with multilevel models; fixed effects: MPM cohort, ROI, 3-way depth interaction term for cubic fit; random effects: subject and ROI [random slope specified for ROI]).

**Figure 2.**
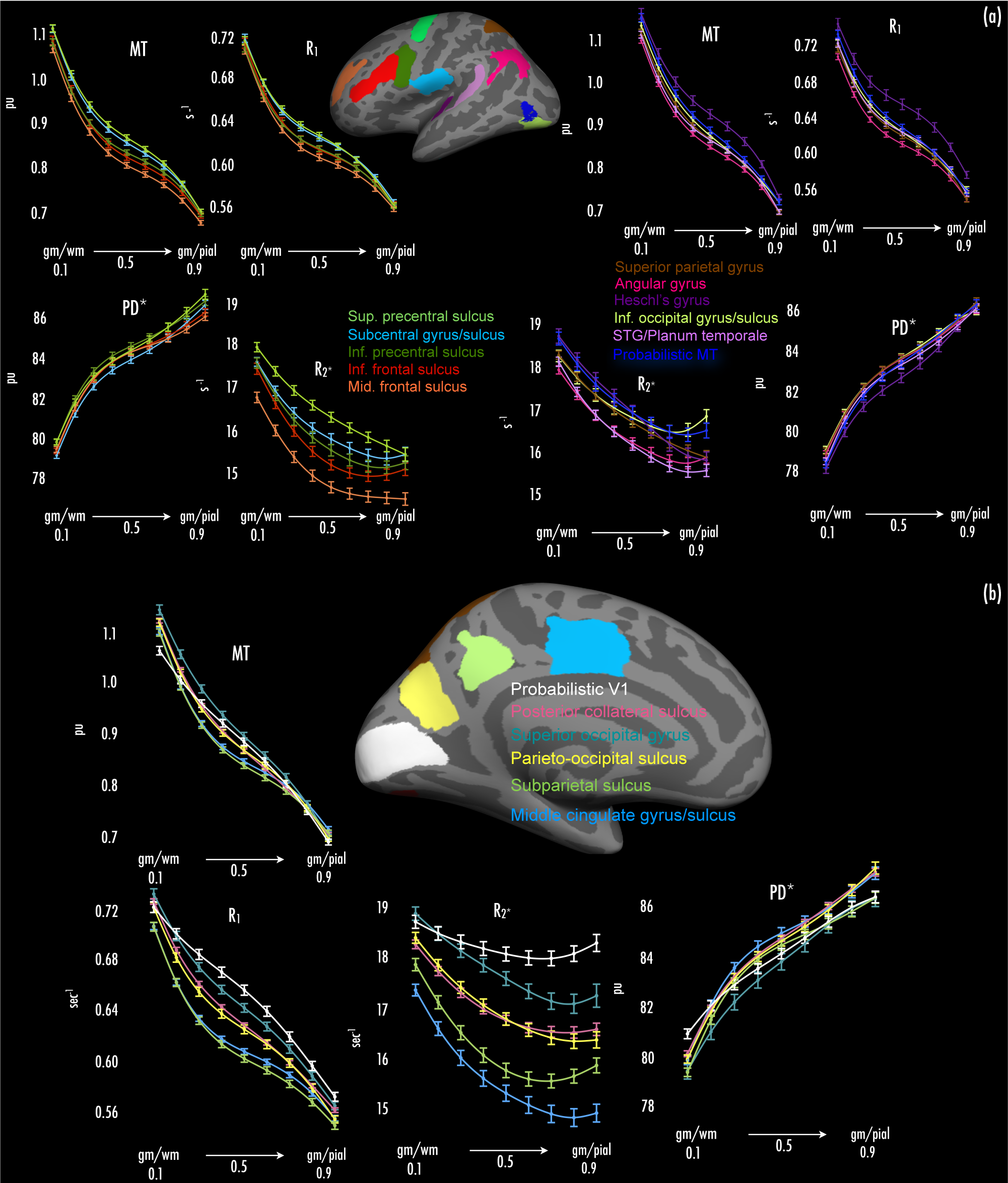
Depth profile plots for multi-parameter maps across cortical ROIs. (a) Depth profiles plots for lateral surface ROIs. Plot panels present mean±95% CI for each parameter across the ROIs (averaging ROI means across hemispheres), extending from proximal to grey-white boundary (gm/wm) to proximal to grey-pial boundary (gm/pial) (x axes, left to right). Line color denotes the ROI (see inset: corresponding ROI color displayed on standard inflated cortical surface; color-coded ROI names listed at centre). (b) Depth profile plots for medial surface ROIs. All other figure specifications as per (a).

R_2_* values also decreased across all ROIs progressing from the white matter to the pial surface. Modeling the reduction in R_2_* values, quadratic trends provided the best fit to data across depth fractions in all ROIs (versus linear and cubic trends – fits specified as for R_1_ and MT, with 2-way depth interaction term for quadratic fit). In many ROIs, R_2_* values showed an upward trend after the initial decrease, particularly at cortical depths close to the pial surface (i.e., 0.8 and 0.9 cortical depth). This pattern was noted in regions including probabilistic MT/V5, inferior occipital gyrus/sulcus, inferior frontal and inferior pre-central sulci, and angular gyrus (see Fig. 2a), as well as probabilistic V1, superior occipital gyrus, and subparietal sulcus (see Fig. 2b). A likely cause of this is the presence of blood vessels close to the pial surface, where high levels of iron within hemoglobin would elevate observed R_2_* values.

PD* values showed increases toward the pial surface from the white matter surface, across all ROIs. Similar to R_1_ and MT values, change in PD* values was greatest at depth fractions proximal to the white matter and pial surfaces (0.1 and 0.9, respectively), with PD* values showing high growth there (cf. the declines noted in R_1_ and MT values). In all ROIs, cubic trends provided the best fit to PD* increases over depth fractions (fits specified as above for R_1_ and MT).

### 3.3 Inter-regional MPM results

Across cortical ROIs, we found clear differences in the mean values of R_1_, MT, R_2_* and PD* parameters, marked by substantial changes in those parameters when sampling across primary and non-primary cortical ROIs.

As expected given previous literature (Glasser et al., 2014; Glasser & Van Essen, 2011; Sereno et al., 2013; Dick et al., 2012; Sigalovsky et al., 2006; Bock et al., 2013; de Martino et al., 2015; Waehnert et al., 2016; Turner, 2015; Nieuwenhuys, 2013), MPMs with highest sensitivity to cortical myelin (R_1_, MT) showed elevated values in ROIs subsuming or proximal to primary cortical areas. These regions included Heschl’s gyrus (Fig. 2a, mauve trace), superior and inferior pre-central sulcus (Fig. 2a, light and dark green traces, respectively), subcentral gyrus/sulcus (Fig. 2a, light blue trace), and probabilistic V1 (Fig. 2b, white trace). R_1_ values in probabilistic V1 were elevated compared to other regions, but note that MT values in probabilistic V1 were not elevated to the same extent seen for R_1_ (see also Fig. 1, MT panels). Other non-primary regions partly characterized by heavier cortical myelin (probabilistic area MT/V5; Walters et al., 2003; Sereno et al., 2013) also had increased R_1_ and MT values (Fig. 2a, dark blue trace).

R_2_*, which typically shows high affinity for tissue iron (a related property of myelinating oligodendrocyte processes; Fukunaga et al., 2010; Todorich et al., 2009), was elevated in most of the regions noted above that were proximal to (or inclusive of) primary cortex and that showed elevated R_1_ and MT values (i.e., Heschl’s gyrus, superior and inferior pre-central sulcus, subcentral gyrus/sulcus and probabilistic V1). As expected, probabilistic area MT also showed elevated R_2_* values. Importantly, surface averaged MPM data (Fig. 1) further suggested that foci of high R_2_* were displaced towards sulcal regions adjacent to foci of high R_1_ and MT, with R_1_ and MT foci manifesting largely at gyri (e.g., compare R_2_* foci at Heschl’s sulcus bilaterally, versus R_1_ and MT foci at medial Hesch’s gyrus, Fig. 1). Further to the account of R_1_ and MT differences above (Group average MPM results, see 3.1), the displacement of R_2_* foci may reflect detection of iron-rich oligodendrocyte cell bodies within sulcal depths (e.g., Stüber et al., 2014); here, thinner cortex relative to adjacent gyri may limit the extent of macromolecule expression, in tandem with the greater signal expressed via oligodendrocytes. Regions of association cortex (e.g., subparietal sulcus, middle cingulate gyrus/sulcus) generally associated with light myelination (Glasser & Van Essen, 2011; Cohen-Adad et al., 2012), showed low R_2_* and correspondingly low R_1_ and MT values (Fig. 2b, light green and light blue traces).

PD* tended to be reduced in heavily-myelinated areas (e.g., Heschl’s gyrus; Fig. 2a, mauve trace). However, we note that inter-regional PD* curves did not reflect a strict ‘mirror-image’ of areas with elevated R_1_ and MT values. For instance, while superior pre-central sulcus (Fig. 2a, light green traces) showed elevated R_1_ and MT curve values, PD* curve values in this region were also increased compared to adjacent cortical areas (see Fig. 2a, PD* panel).

### 3.4 Age effects on R_1_/MT/R_2_*

Exploring development of myelin and related tissue processes cross-sectionally, we correlated age in years with vertex-wise R_1_, MT and R_2_* values that had been residualized by cortical thickness, curvature and MPM cohort. To limit bias in model fits, we calculated ‘leave-one-out’ estimates of models per MPM using a jackknifing procedure (see 2.4.2).

We found evidence of significant (peak-level FDR *q* < 0.05 per hemisphere) correlations between age and R_1_, and age and MT, across the lateral cortical surface. However, we did not find any regions where age and R_2_* correlations survived with FDR-correction (*q* > 0.05).

Positive age-R_1_ correlations (i.e., increasing R_1_ with age; Fig. 3a) were widespread, extending across much of pre-frontal, frontal and parietal cortex bilaterally. These positive age-R_1_ correlations were observed bilaterally at pars opercularis, middle frontal gyrus (MFG), pre-central gyrus and central sulcus, superior frontal gyrus, and superior and inferior parietal lobules (including angular and supramarginal gyri). Additional positive correlations were found in right occipito-temporal regions, including a peak proximal to visual area MT/V5. A further peak emerged at right superior temporal sulcus.

**Figure 3.**
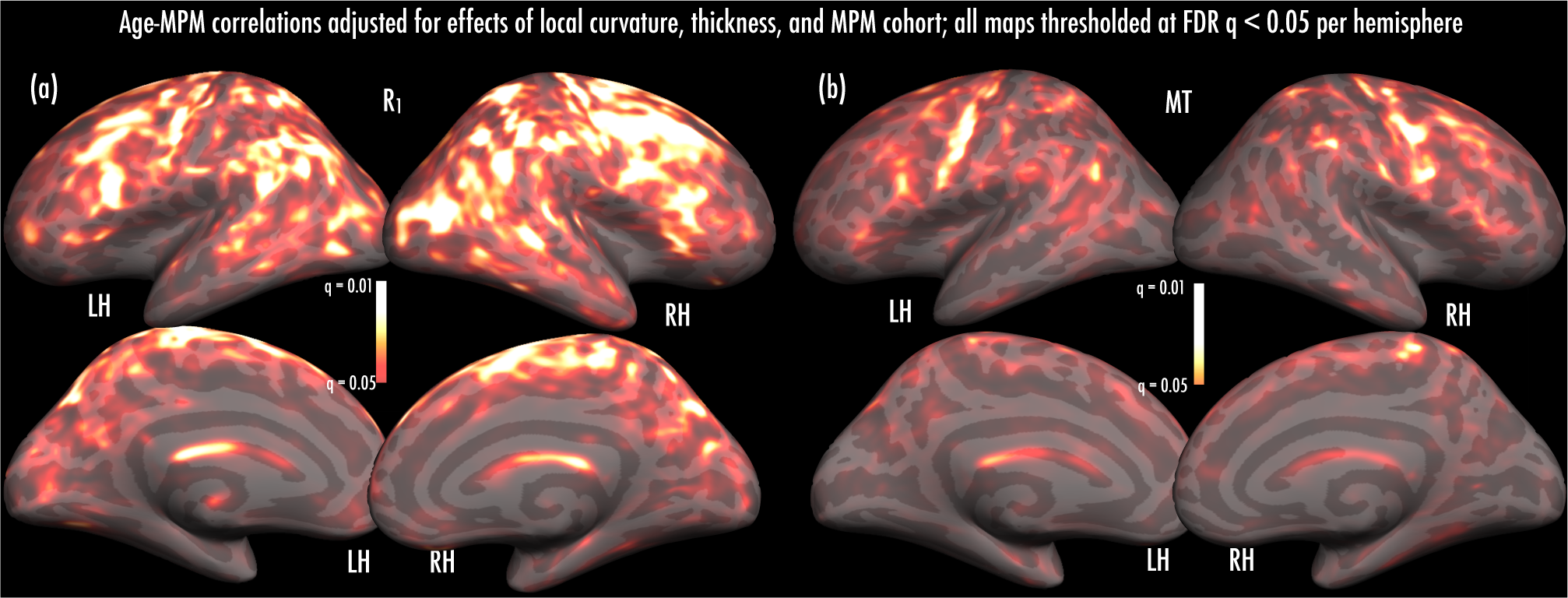
Results of age-MPM correlations. Maps present vertices where positive age-parameter correlations (jackknifed) were significant, after adjusting for effects of local cortical curvature and thickness, and MPM cohort (see Materials and Methods). (a) Significant age-R_1_ correlations emerged across much of the lateral surface, including frontal, parietal and temporal regions. (b) Significant age-MT correlations were less extensive, emerging at pre-central, dorso-lateral pre-frontal, and parietal regions. Heat scale overlays indicate range of FDR-corrected significance of age-parameter correlations (*q* ≤ 0.05-0.01; all effects hemisphere-wise FDR-corrected).

Positive age-MT correlations were less extensive than those observed for age-R_1_ (see Fig. 3b). The largest age-MT peaks manifested at left pre-central gyrus and central sulcus, along with a series of peaks across right pre-central gyrus. Other smaller peaks emerged at right supramarginal gyrus, left pars opercularis, right MFG, and right superior parietal lobule.

### 3.5 Overlap of age effects - R_1_ and MT

As a way to explore the extent to which age effects were common to both R_1_ and MT, we compared age effects for overlapping significant vertices in both analyses. We isolated vertices over each hemisphere where both R_1_ and MT (corrected for covariates) had shown FDR-significant jackknifed age correlations (i.e., *q* < 0.05), and then defined these regions as ROIs, which we sampled onto each subject’s surface (see 2.4.2). Figure 4 presents age (range: 18-39 years) regressed against the ROI mean R_1_ or MT value per subject, over each hemisphere.

**Figure 4.**
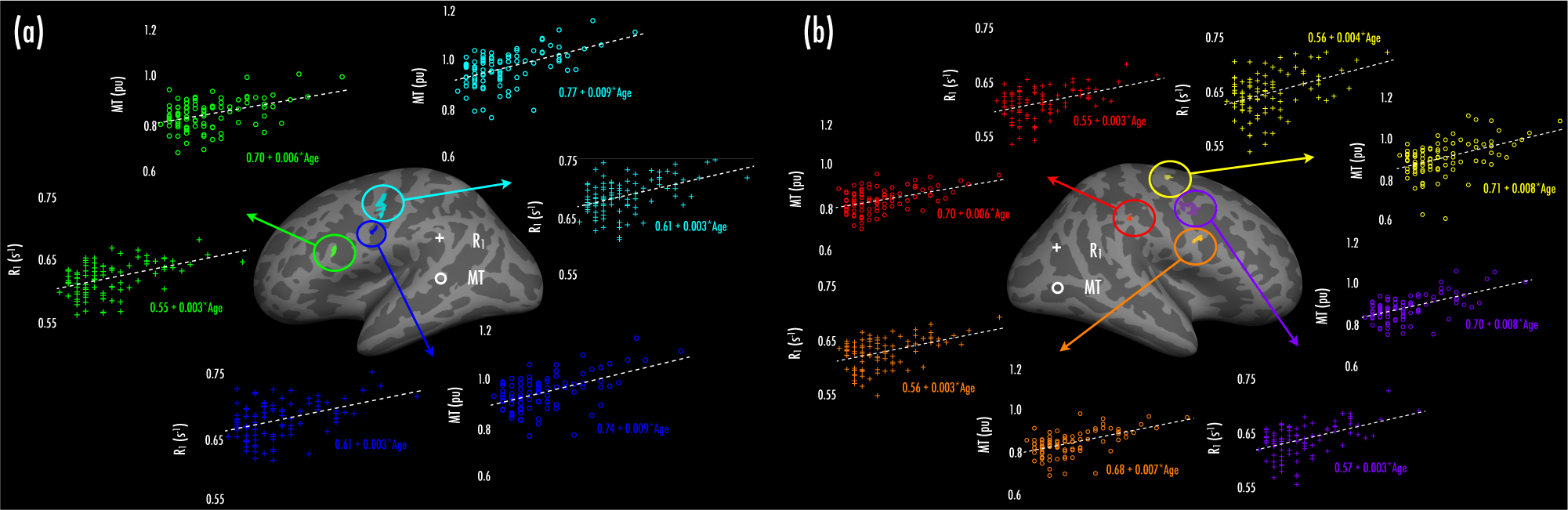
Overlap of R_1_- and MT-age effects over (a) left and (b) right hemispheres. Insets present ROIs, defined as regions where overlap of age-R_1_ and age-MT correlations manifested (at FDR-corrected significance when adjusting for effects of local cortical curvature and thickness, and MPM cohort). Surrounding panels display scatter plots of age and subject-wise ROI mean parameter values (ROI correspondence denoted by color coding), with age-parameter linear fits (dashed traces; intercept and age slope from linear fits reported). Crosses: R_1_; circles: MT.

Across the left hemisphere (Fig. 4a), we found three regions of overlap for age-R1 and age-MT effects; these encompassed pars opercularis (green), lateral pre-central gyrus/central sulcus (cyan), and ventral central sulcus (blue). Linear regression fits for R_1_ showed that across these ROIs, R_1_ values increased at a rate of 0.003 s^-1^ per year. Similarly, over these ROIs, MT values increased at rates ranging from 0.006-0.009 pu per year. Across the right hemisphere (Fig. 4b), we identified four regions where age-R1 and age-MT effects overlapped. These included: dorsal (yellow), lateral (purple), and ventral (orange) pre-central gyrus, and supramarginal gyrus (red). Similar to the left hemisphere, R_1_ values in the right hemisphere ROIs increased at rates of 0.003-0.004 s^-1^ per year; MT values increased at rates ranging from 0.006-0.008 pu per year.

## Discussion

A fundamental challenge in human neuroscience is the development of efficient and robust anatomical imaging techniques that can enable specific tissue properties within the cortex to be quantified *in vivo*. Such approaches are critical to charting healthy human brain structure across development, together with informing understanding of tissue deterioration in aging and disease (Yeatman et al., 2014; Deoni et al., 2015, 2011; Callaghan et al., 2014a; Laule et al., 2004). Here, we used a previously well-validated quantitative multi-parameter mapping (MPM) protocol (Helms et al., 2008a, 2008b; Weiskopf et al., 2011, 2013; Lutti et al., 2011; Callaghan et al., 2014b) to characterize tissue profiles *in-vivo* across a range of cortical regions. Moreover, we explored effects of aging across late adolescence and early adulthood on myelination, using MPM indices that offer tissue specific proxies for cortical myelin processes. We found that across a series of cortical ROIs, MPMs sensitive to myelin and myelin processes (R_1_, MT) showed enhanced values in line with expected differences in myelination between primary and primary adjacent regions, as compared to many non-primary cortical regions. Further, we found that depth profiles of MPMs reflective of myelin and myelin-related processes (R_1_, MT, R_2_*) showed a monotonic decline in parameter values across the cortex, when progressing from the white matter surface towards the pial surface. In contrast, the MPM reflective of tissue water (PD*) increased as a function of distance from the white matter surface. Cross-sectional effects of age on myelination were most robust for R_1_, and manifested as R_1_ increases of ∼0.5% per year across much of pre-frontal, frontal and parietal cortex. Robust aging effects as indexed by MT were change of ∼1.0% per year, but less extensive spatially than those found for R_1_, encompassing pre-central gyrus, along with regions of association cortex.

### 4.1 Inter-regional MPM normative data and intra-regional depth profiles

A central aim of our study was to explore myelin and related tissue processes across a series of cortical regions using MPMs, in order to provide normative mapping data from a healthy sample for these tissue-sensitive metrics. In line with previous histological (Annese et al., 2004) and combined histological/MR investigations (Walters et al., 2003; Stüber et al., 2014; Fracasso et al., 2016), we found that MPMs allowed us to distinguish between cortical regions (e.g., primary vs. association cortex), and also to chart intra-regional tissue properties, based on profiles of tissue-sensitive parameters through the cortical sheet.

Previous investigations have explored inter-regional and cortical depth profiles for R_1_ (=1/T_1_) alone, which has typically shown high sensitivity to heavily-myelinated cortical tissue. In particular, Sigalovsky et al. (2006) charted cortical R_1_ values across anatomical subdivisions of Heschl’s gyrus, while Dick et al. (2012) characterized the cortical depth profile of R_1_ values within divisions of Heschl’s gyrus (Te 1.0, 1.1 & 1.2); both studies found the expected pattern of elevated R_1_ values at the postero-medial aspect of Heschl’s gyrus, reflecting putative auditory core. In visual cortical regions, Sereno et al. (2013) charted the increased R_1_ values found within primary (e.g., V1) and non-primary (e.g., MT, V3a, V6) higher-visual areas, with respect to retinotopic functional borders (see also Abdollahi et al., 2014). Moreover, Sereno et al. demonstrated an overall increase in R_1_ in these visual regions compared to association cortex (angular gyrus), across intra-regional depths.

Here, we re-capitulate many of the findings above, and extend these results to further tissue-sensitive MPMs. Both R_1_ and MT metrics in our present results manifested higher values within primary, primary adjacent, and higher visual cortical regions (e.g., Heschl’s gyrus, probabilistic V1, probabilistic MT/V5, subcentral gyrus) compared to association areas (e.g., middle cingulate sulcus and gyrus, subparietal sulcus). In particular, that our inter- and intra-regional magnetization transfer (MT) results closely mirror our R_1_ data lends strong support to the feasibility of mapping MT as a myelin-proxy in the cortex. A potential benefit here is that MT is less affected by other properties of the tissue microstructure than R_1_ (i.e., R_1_ is partially influenced by R_2_* signal properties, which vary as a function of susceptibility effects due to paramagnetic molecule concentrations – a point we return to below; Callaghan et al., 2014b; Fukunaga et al., 2010; Stüber et al., 2014). MT therefore largely reflects the macromolecular content of the tissue microstructural environment, and can afford a proxy for the bound water fraction (Callaghan et al., 2014b). Moreover, previous post-mortem comparisons of MT ratio and T_1_-based MR metrics have shown high correlation between the two (r=-0.79; Schmierer et al., 2004), suggesting that both converge well toward indexing myelin tissue. A noted difference in R_1_ and MT however was the reduced sensitivity of MT to myelin content within sulcal regions where cortex is thinnest, particularly across the calcarine fissure. As discussed in results and above, the partial R_2_* influence on R_1_ may have increased the contrast-to-noise for myelin and related processes within this sulcal region, where cortical thickness can fall to ∼1mm, reducing the number of voxels that contribute to measured signal. The multiple signal contributions to R_1_ therefore may enhance its sensitivity where measurements of tissue microstructure are required at a particularly fine-grained level, as in very thin and concave cortical regions.

A further consideration in the present results was the convergence between regions that showed high myelin content (i.e., elevated R_1_ and MT values) and also high R_2_* values. The correspondence between R_1_, MT and R_2_* has been charted previously. In particular, tissue molecules such as iron influence R_2_* (Langkammer et al., 2010; Fukunaga et al., 2010), by causing local inhomogeneities in the B_0_ field, in turn influencing the longitudinal relaxation rate (i.e., R_1_) (Callaghan et al., 2014b). Moreover, tissue regions high in iron often reflect areas of high myelin content, since oligodendrocyte cell bodies (whose cytoplasmic membranes extend around axons to form the myelin sheath) are known to express high concentrations of iron (Stüber et al., 2014; Bartzokis, 2004; Todorich et al., 2009). Similarly, regions that show reduced MT in aging (reflecting presumptive demyelination) also tend to manifest reductions in R_2_* (Callaghan et al., 2014a). In line with the inter- and intra-regional variation we observed in R_1_ and MT, R_2_* tended to follow similar profiles of elevation or reduction within regions of respectively high or low R_1_ and MT, mirroring its close relationship with R_1_ and MT. Nevertheless, an important difference emerged in the location of high R_2_* foci, such that these were displaced into sulci adjacent to some of the regions of high R1 and MT (e.g., at Heschl’s sulcus and gyrus, and at infero-temporal sulcus, adjacent to area MT; Fig. 1). The exact mechanisms underlying this difference are unclear; however, a possible account is an interaction between the thinness of cortex over sulcal depths (as compared to thicker and more convex gyri), and the relative expression of macromolecular (i.e., myelin lipid) versus glial content as a result. The limited layer IV/V thickness within deep sulci constrains the expression of high macromolecular content, and in tandem, it is possible that oligodendrocytes may be expressed in sulcal regions proximal to heavily-myelinated gyri (see Stüber et al., 2014). In combination, these differences may lead to apparently elevated R_2_* in sulci adjacent to foci of high R_1_ and MT.

Finally, we observed that intra-regional PD* parameters followed an expected pattern contrary to R_1_, MT, and R_2_*, with highest values at cortical depths close to the pial surface. Agreeing with previous evidence of reduced myelin in superficial cortical layers (Annese et al., 2004; Walters et al., 2003; Leuze et al., 2014), the high PD* values likely reflect increased tissue water, owing to the high grey matter volume, dense vascularization and low myelin lipid content at the pial surface. The observed changes are in line with those that use 1 minus the ratio of PD and cerebrospinal fluid as a marker for macromolecular tissue volume (Mezer et al., 2013).

### 4.2 Development and cortical myelination

Here, we found evidence of a protracted course of myelin and myelin-related process development within the cortex, across late adolescence and early adulthood. Our MPMs varied in the extent to which they revealed these developmental effects to be robust (i.e., to hemisphere-wise FDR correction, *q* < 0.05): R_1_ showed widespread increases with age across pre-frontal, frontal and parietal cortex; MT revealed a slightly lesser extent of age effects that were statistically significant with multiple comparison correction, largely concentrated in pre-central regions bilaterally.

Previous MR studies have shown that myelination in humans begins in the subcortex (Partridge et al., 2004; Deoni et al., 2011; Barkovich et al., 1988; Nakagawa et al., 1998; for review, see Paus et al., 2001; Baumann & Pham-Dinh, 2001). Further, MR studies have found myelination of primary and association cortex progresses during childhood (Deoni et al., 2015; Dean et al., 2016) and adolescence (Grydeland et al., 2013), before age-related de-myelination begins during middle adulthood (Vidal-Piñeiro et al., 2016; Grydeland et al., 2013; Salat et al., 2009; see also Rowley et al., 2017). DTI investigations have also found that white matter structure within association cortex develops over extended periods, typically to beyond late adolescence (Klinberg et al., 1999; Barnea-Goraly, 2005). Quantitative assays of subcortical fibre myelination with R_1_ have similarly shown an inverted-U profile, indicative of protracted myelin development during childhood, adolescence and early adulthood, followed by de-myelination from middle to older age (Yeatman et al., 2014).

Of particular relevance to our present results, recent studies using T_1w_/T_2w_ image contrast ratios have identified extended periods of white matter development within association cortex. Shafee et al. (2015) and Grydeland et al. (2013) found significant increases in T_1w_/T_2w_ ratio with age over much of the frontal and parietal lobes, in young adults (18-35 years) and across the lifespan (8-83 years), respectively. Notably, the linear trend identified in the present study (and by Shafee et al.) was also found by Grydeland et al. when considering the younger (8-20 year old) tail of their age distribution (cf. quadratic trends in T_1w_/T_2w_ ratio across their full age range). This appears to reflect phases of increasing myelination up to early adulthood, followed by periods of relative stability and eventual decline of cortical myelin after 50-60 years of age (Grydeland et al., 2013; see also Miller et al., 2012).

As outlined above, an important advance made by our age findings is our use of MPMs that index a range of myelin-related tissue processes. MPMs enabled us to and paramagnetic ions (R_1_), together with either predominantly macromolecular (MT), or paramagnetic ion (i.e., iron) (R_2_*) concentration (Callaghan et al., 2014b). Existing studies that have used T_1w_/T_2w_ ratio methods cannot resolve for tissue-sensitive parameters, since both T_1w_ and T_2_w contrast are determined by a variety of microstructural (see Glasser et al., 2014; Vidal-Piñeiro et al., 2016), and vasodilatory properties (i.e., via O_2_/CO_2_ concentration; Tardif et al., 2017). Our finding of regions that manifested overlapping age effects for R_1_ and MT suggests that myelination follows a protracted developmental course. Interestingly, we did not observe robust age effects on R_2_*, a parameter sensitive to tissue iron concentration. Whereas previous studies have shown focal R_2_* decreases along fibre bundles (cf. sub-cortex) in line with aging in older samples (Callaghan et al., 2014a), here, our younger cohort showed increases in parameters largely reflective of cortical myelin content. One speculative account is that the developmental effects we observed involved changes to myelin sheath thickness (i.e., g-ratio; Dean et al., 2016) rather than large-scale changes in the numbers or density of iron-rich oligodendrocyte cell bodies. However, optogenetic evidence in mice has supported effects of behaviorally-relevant neural activity in promoting increases in both myelin sheath thickness (i.e., g-ratio decreases) and oligodendrogenesis (Gibson et al., 2014). Taken together, it is likely that our current effects of age reflect some combination of these processes, although the precise mechanisms remain unclear. Future studies in which *in-vivo* measurements of g-ratio (e.g., Mohammadi et al., 2015) are probed across our age range in addition to each multi-parameter map may enable us to shed further light on the mechanisms at play in myelin sheath development.

A strength of the present approach is the use of a well-validated MPM protocol incorporating B_0_ and B_1_ RF transmit field mapping. Here, field maps form an integral part of the MPM protocol, such that local flip angles can be resolved and used in the estimation of MPMs, based on a variable flip angle procedure (see Lutti et al., 2010, 2012; Helms et al., 2008b; Weiskopf et al., 2011). T_1w_/T_2w_ ratio methods are subject to B_1_ RF transmit field inhomogeneities during acquisition, which can bias local flip angles and the resulting signal intensity and contrast, increasing measurement error (Lutti et al., 2010; Helms et al., 2008b; Weiskopf et al., 2011; Lutti et al., 2014; but see also Glasser et al., 2014). The precision and reproducibility of the MPM and field mapping protocols have been documented previously (Weiskopf et al., 2013; Lutti et al., 2010, 2012). Importantly, although we observed differences in MPM parameter values between the protocols that differed across cohorts in our present study, those differences reflected a constant offset, the source of which was isolated and which we controlled for in statistical models.

In light of these methodological advances, our present findings agree well with accounts of cortical development and myelination based on T_1w_/T_2w_ ratio methods. Moreover, that we were able to identify robust linear developmental effects on cortical myelination/myelin processes and with a much smaller cohort than many studies (Shafee et al., 2015; Grydeland et al., 2013) suggests that reliable quantitative mapping protocols can play a highly informative role in charting cortical development.

### 4.3 Conclusions

Using a well-validated multi-parameter mapping protocol, we showed that both inter- and intra-regional cortical myelin and related processes can be quantified across much of the cortical sheet. We further demonstrated the utility of delineating profiles of cortical myelin processes across cortical depths (using R_1_, MT, and R_2_*) in tandem with mapping of tissue water sensitive parameters (PD*). Moreover, exploring effects of development cross-sectionally, we found that cortical myelin and myelin processes increased at a rate of 0.5 % (R_1_) per year over frontal and parietal regions, across late adolescence and early adulthood. These results shed further light on ontogenetic factors that may shape large-scale cortical organization and inform broader accounts of lifespan cortical development, as well as helping to characterize the healthy aging of the human brain, which may provide a useful clinical benchmark for studying de-myelination in aging and disease.

## Acknowledgements

We thank Martin Sereno for many custom changes to csurf that facilitated this work.

## Supplemental Methods

### Image pre-processing and surface reconstruction

#### Image synthesis

First, two synthetic FLASH volumes were created using the FreeSurfer mri_synthesize routine. Inputs to the routine were scaled quantitative PD and T_1_ (1/R_1_ volumes, with removal of a small number of negative and very high values produced by estimation errors. For the first synthetic image, default FreeSurfer contrast parameters were specified. The second synthetic image was produced using the same PD and T_1_ input volumes, with contrast parameters specified (TR = 20 ms; α = 30°; TE = 2.5 ms). During synthesis, images were ‘conformed’ to 1mm^3^ isotropic resolution in FreeSurfer. Both synthetic images were then further scaled with AFNI 3dcalc; this additional linear scaling yielded image intensity properties closer to the optimal intensity values needed to segment tissue boundaries in FreeSurfer. The image synthesized with default contrast parameters was used as the main input to the FreeSurfer automated processing stream (following further pre-processing steps; see below). The image synthesized with specified contrast parameters was used at a later stage as input to the FreeSurfer Talairach transformation. Finally, a scaled and truncated version of the PD volume was produced with AFNI 3dcalc. This adjusted PD volume was used as input to the skull strip procedure (see below).

#### Manual image adjustment

Because the FreeSurfer reconstruction pipeline relies on highly homogeneous values within tissues, the FLASH image synthesized with default parameters was further adjusted for subjects within the first cohort (*n* = 34) using an inhouse version of FreeSurfer (Csurf). Differences in the MPM acquisition led to improved image contrast in the second cohort (*n* = 59), and thus manual image adjustment was not required. Each subject’s synthetic image was hand-adjusted using a piecewise linear normalization procedure to linearly ramp intensity values of grey and white matter within isolated regions. Brightness values of voxels within the inferior and medial temporal lobes, temporal pole, long and short insular gyri, and ventro-medial pre-frontal cortex were gently rescaled (< 1.2x). Manual blink comparison between the synthetic volume and the labelled white matter surface was used to compare adjustments as each brightening iteration was applied. Care was taken to ensure that manual brightening did not cause grey and white matter to exceed the intensity value bounds specified for those tissue classes in FreeSurfer (grey matter: 50-70; white matter: 100-140). Manually brightened synthetic images were saved and used within the skull strip procedure.

#### Skull strip

Next, the subject’s adjusted quantitative PD volume (see image synthesis) was used as input to a customized skull strip procedure run in Csurf. Briefly, the skull strip procedure removed the skull and regions exterior to it from the image volume, rendering an image of remaining brain tissue (including cerebellum and brainstem). First, an elliptical surface (4th or 5th geodesic subtessellation of an icosahedron) was expanded from inside the PD volume, with expansion of the surface constrained by arrival at low intensity voxels (i.e., those containing CSF and/or the inner surface of the skull). The set of voxels intersecting the faces of the resulting surface was then flood-filled from the outside, thereby constraining the brain volume to the brighter voxels inside the surface region. Using this PD volume as a mask, flood-filled voxels in the volume were used to set the corresponding voxels in the subject’s default-parameter synthetic image to an intensity of zero. The boundaries of the flood-filled voxels within the skull-stripped PD image were then manually adjusted to correct for any local deviations into neural tissue (particularly in regions proximal to paranasal sinuses, prone to susceptibility artifacts). Manual adjustment involved reducing the intensity threshold for cortical grey matter (to a value of 40); the flood-filled boundary was then forced toward voxels below this threshold. Manual adjustment was applied to the synthetic volume; the skull-stripped synthetic volume was used as input to a custom version of the surface reconstruction pipeline.

#### Surface reconstruction

First, each subject’s skull-stripped synthetic volume was intensity normalized in FreeSurfer (using the mri_normalize routine). Normalized images were briefly inspected to ensure grey and white matter intensity values were within the appropriate ranges (white matter: 110; grey matter: 50-70). Next, the skull-stripped default parameter synthetic volume (see skull strip) was used to mask the contrast-specified synthetic volume (see image synthesis). This masked (i.e., skull-stripped), contrast- specified synthetic volume was then used as the input volume to an initial Talairach transformation process (run using the FreeSurfer mri_em_register routine). Next, a further normalization step was performed (using the -canorm parameter in FreeSurfer recon-all); the initial Talairach transform, the skull-stripped, default parameter synthetic volume and the intensity-normalized version of that volume were used as inputs. Following this, a multi-dimensional Talairach transformation was applied (using the -careg and -careginv parameters in recon-all); the normalized volume (produced by recon-all -canorm), the skull-stripped default parameter synthetic volume, and the initial Talairach transform were used as inputs. Finally, the full FreeSurfer recon-all pipeline was run for each subject (parameters specified can be found at: https://surfer.nmr.mgh.harvard.edu/fswiki/ReconAllDevTable; parameters used were those of the autorecon-2 stage, and the first 6 parameters from autorecon-3).

## Footnotes

1. For the first MPM cohort (*n* = 34), T_1w_, PD_w_ and MT images were acquired with different slab orientations over some subjects relative to others. Initial inspection of data acquired with the slab aligned to each cardinal axis showed susceptibility artifact that affected cortex in a subset of participants. Although eye movements were monitored during scanning runs, slight movement (e.g., due to blinking) led to artifact within orbitofrontal and medial temporal lobes in some datasets. To counter this issue, the acquisition protocol was modified, by rotating each image slab at 30° about the x-axis (such that the eyes lay outside the slab). Participants with data acquired without slab rotation were inspected blind to subject identity for evidence of susceptibility artifact; those participants that showed evidence of artifact within cortical areas were re-scanned using the rotated acquisition protocol. In total, 6 of 34 participants showed susceptibility artifact with the unrotated acquisition and were re-scanned with the rotated protocol; 15 of 34 participants showed no evidence of susceptibility artifact with the original unrotated acquisition and were not re-scanned; 13 of 34 participants were scanned using the rotated protocol as default. Whole-brain analyses of each MPM using slab rotation as a regressor of interest showed no significant differences across any vertices over either hemisphere for participants with rotated versus unrotated acquisition (*p* < 0.01, uncorrected). Moreover, there was no significant difference in age between the subjects with non-rotated versus rotated acquisition (z < 0.01, *p* > 0.99).

## Notes

The research leading to these results received funding from the European Research Council under the European Union's Seventh Framework Programme (FP7/2007-2013) / ERC grant agreement n° 616905, and via EC FP7 grant n° MC-ITN-264301 (TRACKDEV) to DC. The Wellcome Trust Centre for Neuroimaging is supported by core funding from the Wellcome Trust 0915/Z/10/Z.

